# Linking micro and macroevolution in the presence of migration

**DOI:** 10.1101/490128

**Authors:** Pablo Duchen, Sophie Hautphenne, Laurent Lehmann, Nicolas Salamin

## Abstract

1. The process of speciation is of key importance in evolutionary biology because it shapes macroevolutionary patterns. This process starts at the microevolutionary level, for instance, when two subpopulations evolve towards different phenotypic optima. The speed at which these optima are reached is controlled by the degree of stabilising selection, which pushes a mean trait towards an optimum within subpopulations, and ongoing migration that pulls the mean phenotype away from that optimum. Traditionally, macro phenotypic evolution with selection has been modelled by Ornstein-Uhlenbeck (OU) processes, but these models have ignored the role of migration within species.

2. Here, our goal is to reconcile the processes of micro and macroevolution by modelling migration during speciation. More precisely, we introduce an OU model where migration happens between two subpopulations within a branch of a phylogeny and this migration decreases over time as it happens during speciation. We then use this model to study the evolution of trait means along a phylogeny, as well as the way phenotypic disparity between species changes with successive epochs.

3. We show that ignoring the effect of migration in sampled time-series data leads to a significant underestimation of the selective forces acting upon it. We also show that migration decreases the expected phenotypic disparity between species and we show the effect of migration in the particular case of niche filling. We further introduce a method to jointly estimate selection and migration from time-series data.

4. Our model extends standard results of interactions selection-migration in a microevolutionary time frame across multiple speciation events at a macroevolutionary scale. Our results further proof that not accounting for gene flow has important consequences in inferences at both the micro and macroevolutionary scale.

## 1 Introduction

The study of macroevolution has proven useful in addressing key evolutionary questions about the build-up of biodiversity and the mechanisms underlying the speciation process (Stanley 1979; Lande 1980b; Futuyma & Agrawal 2009; Katzourakis *et al.* 2009; Campbell & Kessler 2013). These questions have been addressed by modelling, across a phylogeny, the changes in the rate of evolution of a phenotype (e.g. O’Meara *et al.* 2006; Slater *et al.* 2012), the rate of diversification of species (e.g. Simpson 1944; Nee *et al.* 1992; Jablonski 2008; Silvestro *et al.* 2011; Stadler 2011; Morlon 2014), or the effect of a trait on species diversification (e.g. Rieseberg *et al.* 2002; Cardillo *et al.* 2005; Clauset & Erwin 2008; FitzJohn 2012). Although applications of macroevolutionary models are firmly grounded in evolutionary biology (Simpson 1953), the recent theoretical developments in modelling macroevolution have helped understand the mechanisms underlying phenotypic changes across lineages (e.g. FitzJohn 2010; Landis *et al.* 2012).

One of the earliest and main focus of macroevolution has been testing hypotheses about the evolution of quantitative traits among related species (Felsenstein 1985; 2004). Along these lines, neutral trait evolution has been the standard null model for most macroevolutionary studies, and this is typically modelled by Brownian motion (BM). However, the need to incorporate biologically relevant features (e.g. Hansen 1997; Uyeda *et al.* 2011) has lead to large methodological developments (Edwards *et al.* 1964; Cavalli-Sforza & Edwards 1967; Hansen & Martins 1996; Freckleton 2012; Brawand *et al.* 2011; Duchen *et al.* 2017; Boucher *et al.* 2017). One such relevant feature is natural selection, which, in its simplest form acts as stabilising, with traits being pushed towards an optimum. In the presence of stochastic effects on phenotypic change, stabilising natural selection can sometimes be modelled with an Ornstein-Uhlenbeck (OU) process (e.g. Lande 1976, p. 324), which entails a linear transformation of the phenotype making the analysis generally tractable (Gardiner 2009). Since then, the OU process has been standard when representing stabilising selection in models of macroevolution (Felsenstein 1988; Hansen & Martins 1996; Cooper *et al.* 2016).

However, variation in phenotypic data at the macro scale is often difficult to be explained with just a one dimensional OU process representing directional selection (Pennell *et al.* 2015), and if multi-species datasets are small OU tends to be incorrectly favoured over simpler scenarios (Cooper *et al.* 2016). The application of an OU process in macroevolution therefore requires further developments and finer scrutiny. New theoretical developments should thus start from microevolutionary dynamics, and, from this, try to derive macroevolutionary dynamics. A theoretical description of current macroevolutionary models showed that inter-specific trait-covariances depend on microevolutionary forces, such as random genetic drift, stabilising selection, and mutation, at each generation (Hansen & Martins 1996).

The model of Hansen & Martins (1996) nevertheless overlooked the potential role that migration or gene flow within species plays in linking micro and macroevolutionary dynamics, and thus a more detailed connection between these processes is still needed (Salamin *et al.* 2010; Rolland *et al.* 2018). For instance, migration is determinant in setting the speed of divergence between populations, which, in turn, sets the pace at which speciation takes place (e.g. Gavrilets 2004). And more generally, microevolution is fundamentally affected by the interaction between selection and migration in populations subject to limited dispersal (e.g. Wright 1931; Hartl *et al.* 1997; Ronce & Kirkpatrick 2001; Barton *et al.* 2007). Hence, there is a need to study within-species migration when modelling macroevolution to understand the effects of migration on speciation and its interaction with selection.

In this paper, our goal is to connect the processes of micro and macroevolution by modelling migration during speciation. Building on the stabilising selection models of Lande (1976) and Ronce & Kirkpatrick (2001), we introduced a model of phenotypic evolution where migration occurs between two subpopulations before speciation takes place. This model takes the form of an OU process and our approach differs from Bartoszek *et al.* (2017), who modelled migration between branches on a phylogeny, not within as proposed here. Our model lets the trait evolve along the branches of a phylogeny with migration functions decreasing the rate of gene flow through time until speciation happens. We use this model to study the evolution of trait means and phenotypic disparity (with special emphasis on niche filling) between species on a macroevolutionary time scale. Finally, we analysed the effect of migration on the parameter estimates of selection by developing an estimator of the selection coefficient for cases when migration is present or absent and we assess its accuracy with simulations. We show that not accounting for migration can drastically affect the estimation of selection in micro and macroevolutionary models and our approach opens new avenues to better incorporate microevolutionary forces in macroevolutionary modelling.

## 2 Material and Methods

### 2.1 Biological model

Our aim is to model the evolution of a single quantitative phenotype *z* along a phylogenetic tree. Such phylogenetic tree will consist of several epochs, where one epoch is defined as the time span between two successive nodes (see Supplementary Information (SI) Fig. SI-H.1). We first describe the model for one epoch, where microevolutionary forces can change the mean phenotype. We then extend the model to multiple epochs and derive expressions for the expectation and variance of the mean phenotype in each species at the end of each epoch.

#### 2.1.1 One epoch

We assume that each epoch is of length *T* and that the population forming a species in any epoch is divided into two subpopulations of equal and constant sizes.

##### Microevolutionary time scale

For *i* = 1, 2, let 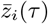 denote the mean phenotype in subpopulation *i*, at time *τ* with initial phenotype 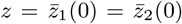, where *z* is a normally distributed random variable with mean *μ* and variance *σ*^2^. The phenotypic evolution of the two subpopulations forming one species is assumed to be characterized by the system of stochastic differential equations

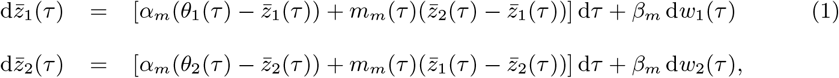

where *θ*_1_(*τ*) and *θ*_2_(*τ*) represent the time-dependent phenotypic optima (or the *phenotypic value* targeted by selection) in each subpopulation at time *τ* (where a time unit is a generation), *α*_*m*_ is the product of the additive genetic variance *σ*^2^ and the strength of selection *γ* on the phenotype, i.e., *α*_*m*_ = *γσ* ^2^ (Lande 1979; Hansen & Martins 1996), and *m*_*m*_(*τ*) is the rate of a migration of an individual at time *τ ϵ* [0, *T*]. Additionally, *w*_1_(*τ*), *w*_2_(*τ*) are two independent Wiener processes with variance 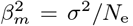, where *N*_e_ is the effective population size. As such, Eq. (1) combines elements of the quantitative genetics models of Lande (1980a, Eq. (3) & (15)) and Ronce & Kirkpatrick (2001, Eq. (2a)), and adds time dependence to the phenotypic optima and the migration rate. From a stochastic process point of view, Eq. (1) is an OU process (Gardiner 2009) and for a single isolated population this model is equivalent to that of (Hansen & Martins 1996). The optima and migration functions characterize the environment of the focal species and we assume that their dynamics are given by

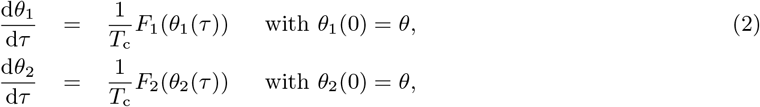

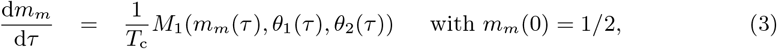

and where *T*_c_ is a characteristic time scale over which the optima and migration rate change in each subpopulation.

Equations (2) describe the change of these optima in each subpopulation as a consequence of environmental change. The characteristic time *T*_c_ will take the value *T*_c_ = 1 when environmental change occurs on the same time scale as the change in phenotype, whereas the optimum changes at a slower rate than the phenotype if *T*_c_ ≫ 1. The *m*_*m*_(*τ*) function describes the migration rate between the two subpopulations. Eq. (3) describes the change of migration over time between populations on their way to speciation. We assume that the time scale over which migration changes is the same as that of the optimum functions.

##### Macroevolutionary time scale

We now change the time scale of the evolutionary process (Eq. (1) and Eq. (2)–(3)) to reach a slower, macroevolutionary time scale *t* defined as

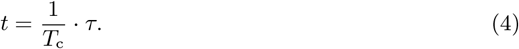

The rescaling is done with the chain rule 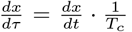, where *x* represents either 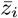, *w*_*i*_, or *θ*_*i*_. The parameters of Eq. (1) and Eqs. (2)–(3) will also be re-scaled such that *α* = *T*_c_*α*_*m*_, *β* = *T*_c_*β*_*m*_, and *m*(*t*) = *T*_c_*m*_*m*_(*t*) to obtain the system of equations

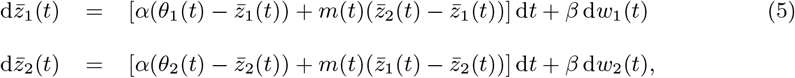

with corresponding phenotypic optima and migration function

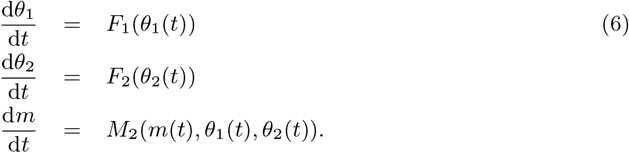

Note that *α* in Eq. (5) accumulates the net effect of phenotypic change due to selection over multiple generations and can thus be interpreted as a macroevolutionary selection coefficient. Finally, if we assume that, at the microevolutionary time scale, selection is weak and that there is a constant but small input of mutation, the genetic variance can be held at its mutation-drift equilibrium and 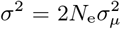, where 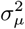 is the mutation variance (Lande 1980a; Hansen & Martins 1996; Walsh & Lynch 2018). Then,

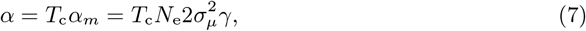

and the variance of the Wiener process is equal to

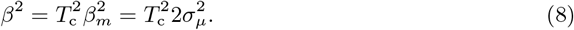

##### Dynamics of the environment

From here on we stay only within the macroevolutionary scale, and we will refer to the macroevolutionary selection coefficient *α* simply as the selection coefficient. We assume that there is random mixing between the two subpopulations at the beginning of each epoch. Over time, migration will decrease and, thus, contribute to population divergence. More specifically, the migration between the two subpopulations follows a monotonically decreasing *migration rate function m*(*t*) (*t ϵ* [0, *T*]) such that *m*(0) = 1/2 (total random mixing), and lim_*t*→∞_ *m*(*t*) = 0. There is a *speciation event* at time *t* if *m*(*t*) < *ϵ*, for a chosen small value *ϵ* > 0 (i.e. migration becomes negligible and *t* = *T*). Throughout the paper we choose *ϵ* = 10^−4^.

We assume that the optima in the two subpopulations are initially the same, *θ*_1_(0) = *θ*_2_(0), but then diverge according to the *differentiation* function *d*(*t*): = *|θ*_1_(*t*) *− θ*_2_(*t*)*|*. As a concrete application of our model, we consider two simple forms of the dynamics of the optima given by 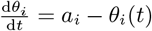 and 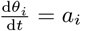. For the initial condition *θ*_*i*_(0) = *θ* (for some initial value *θ*), the solution to these dynamics are given by the following parametric functions *θ*_*i*_(*t*):

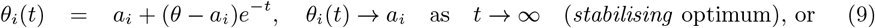

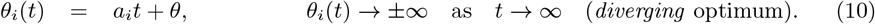

We consider migration functions of the following two types:

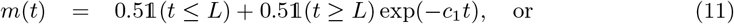

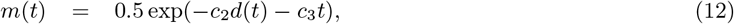

where 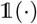 is the indicator function, for some constant parameters *L, c*_1_, *c*_2_ and *c*_3_. Parameter *L* in (11) controls the length of the period during which there is total mixing between the two subpopulations before migration starts decreasing exponentially at rate *c*_1_. In (12), the differentiation function *d*(*t*) affects the decrease rate of the migration function. We present various possible trajectories of 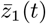 and 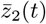 before the first speciation event, with a stabilising optimum in subpopulation 1 and a diverging optimum in subpopulation 2, for different values of *α*, *β*, and *m*(*t*) (Table 1, Fig. 1).

**Figure 1:**
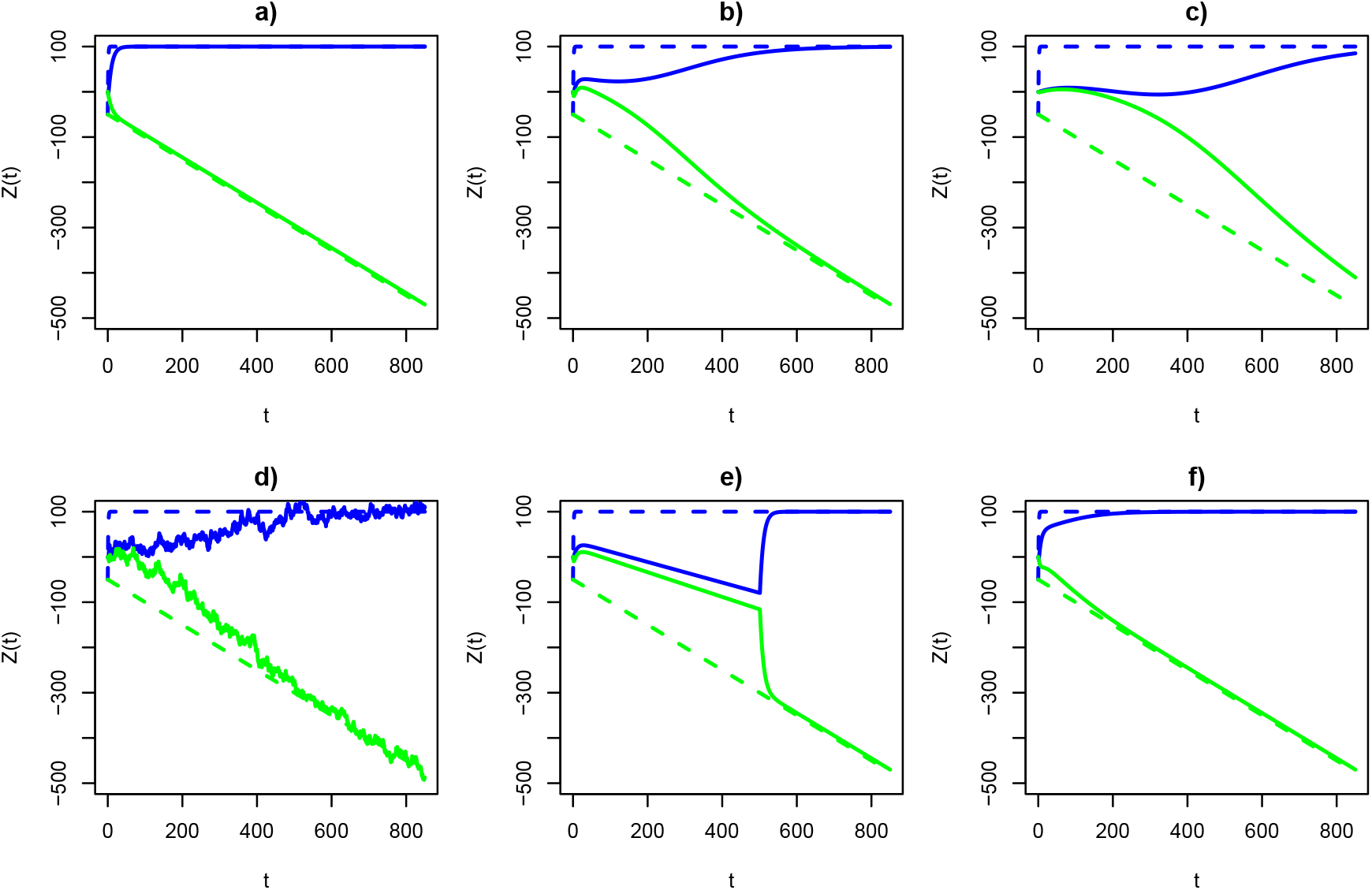
Behaviour of 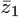 (blue) and 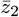 (green) over time *t* under different values of the selection coefficient *α* and the migration parameter *c*_*x*_ (see Table 1 for details in each panel). Actual values of 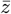 are depicted with solid lines, whereas optima are displayed with dashed lines. These results are discussed in section 3.1.

**Table 1:** Example parameter combinations of the model: selection coefficient *α*, standard deviation of the Wiener process *β*, migration parameters *c*_1_, *c*_2_, *c*_3_, and time span *L* during which there is total mixing (Eq. (11) and (12)). Lastly, we have the biological scenarios associated to each parameter combination, and their corresponding panel in Fig. 1.

| $\alpha$ | $\beta$ | $c_1$ | $c_2$ | $c_3$ | $L$ | Biological scenario | Panel in Fig. 1 |
| --- | --- | --- | --- | --- | --- | --- | --- |
| 0.1 | 0 | - | - | - | - | OU, no migration, no noise | a) |
| 0.1 | 0 | - | - | 0.01 | - | OU with migration, no noise | b) |
| 0.01 | 0 | - | - | 0.01 | - | As above plus weak selection | c) |
| 0.1 | 5 | - | - | 0.01 | - | OU with migration and noise | d) |
| 0.1 | 0 | 0.025 | - | - | 500 | OU with migration, $L > 0$, no noise | e) |
| 0.1 | 0 | - | 0.015 | 0.01 | - | Migration depending on $d(t)$, no noise | f) |

#### 2.1.2 Multiple epochs

We now consider phenotypic dynamics over multiple epochs. To deal with this, the optimum functions can be different for different species and epochs, but we hold the migration functions constant (an alternative scenario, where migration can be different for different species is shown in SI-C.2). The fact that the migration function is fixed implies that the speciation times are deterministic, and so is the number of branches in the phylogenetic tree at any given time: if one initially starts with a single species, then there are 2^*n−*1^ coexisting species during epoch *n*, corresponding to 2^*n*^ subpopulations (*n* ≥ 0).

We denote by 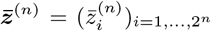 the random vector recording the *mean phenotype of each species* at the end of the *n*th epoch, and by 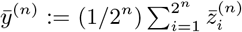 the scalar random variable recording the *averaged mean phenotype* at the end of the *n*th epoch (*n* ≥ 1). We show in SI-C that 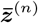 follows a multivariate normal distribution 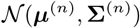 whose mean vector *μ*^(*n*)^ and covariance matrix Σ^(*n*)^ of size 2^*n*^ satisfy a first order recurrence equation,

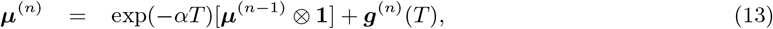

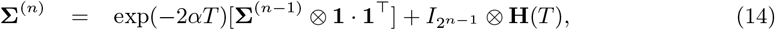

for *n* ≥ 1, with *μ*^(0)^ = *μ* and Σ^(0)^ = *σ*^2^, and where *g*^(*n*)^(*T*) and **H**(*T*) are respectively given by Eq. (50) and Eq. (45); see Proposition C.1. Here, *g*^(*n*)^(*T*) is a sequence of vectors that depend on the optimum functions *θ*_*i*_(*t*), while **H**(*T*) is a matrix that takes into account the covariance induced by the Brownian noises acting on the mean phenotypes of the two subpopulations, and the mass exchange between these subpopulations when *m*(*t*) *> ϵ*.

The vector 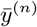 follows a univariate normal distribution with mean and variance given in Eq. (58) and Eq. (59) (see SI-C.1.2). As we further show in SI-C.2, the formulas for *μ*^(*n*)^ and Σ^(*n*)^ can be extended to the case where the migration function *m*(*t*) is different for each species, leading to branches of different lengths in the phylogenetic tree.

Finally, we note that an important descriptor of the phenotypic joint distribution is the *disparity D*^(*n*)^ *of* 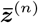 (Harmon *et al.* 2003), which is a scalar random variable measuring the extent to which the mean phenotypes of the species present at the end of the *n*th epoch differ from each other. We define disparity as

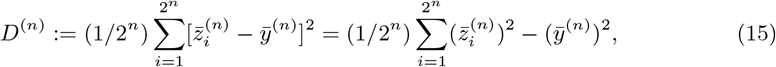

and show in SI-C.1.3 that the first moment of *D*^(*n*)^ is

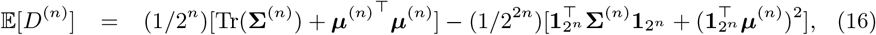

where Tr(Σ^(*n*)^) denotes the trace of the covariance matrix Σ^(*n*)^. Hence, we can evaluate the disparity in terms of Eq. (13)–(14).

### 2.2 Estimation of the selection coefficient and the migration parameter *c*

The applications of OU processes in macroevolution often aim at quantifying the amount of selection experienced by different species without considering the effects of migration. We generalize this to the case with migration and formulate estimators of *α* and the migration parameter *c* = *c*_1_ in Eq. (11) when *L* = 0, that is *m*(*t*) = 0.5 exp (*−ct*). Our model readily lends itself to derive such estimators by setting *β* = 0 in Eq. (5), approximating these expressions as difference equations, and iterating this process *n* times to obtain

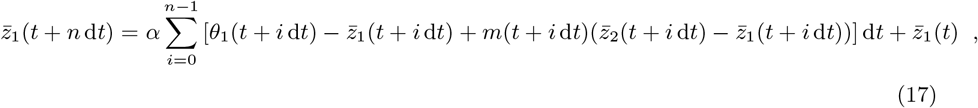

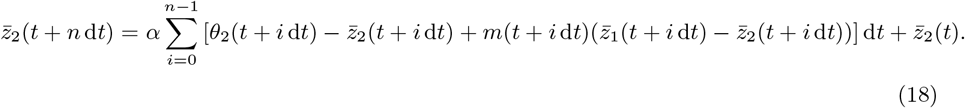

By rearranging terms, the estimator of *α*, denoted 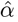, can be written as

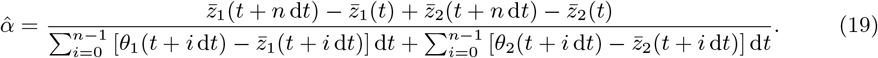

A full step-by-step derivation of Eq. (19) is shown in section SI-F. To obtain an estimator for the migration paramter *c* we simply replace *α* in (17) or (18) with the value of (19) and solve numerically for *c*. If one has data only from a single isolated subpopulation, then *α* can be estimated from Eq. (19) using only the corresponding subpopulation, say subpopulation 1 to estimate *α*:

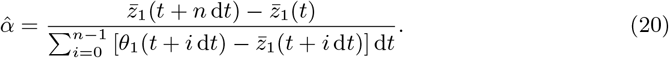

### 2.3 Case study: niche filling

An interesting application of the joint phenotypic distribution and disparity across epochs concerns niche filling. Ecologically speaking, niche filling is a phenomenon by which different populations or species “fill” the phenotypic space of a niche under two conditions: 1) the range of values a phenotype can take is bounded, and 2) two phenotypes cannot take on the same value. This happens, for instance, when there is ecological competition for resources, which prevents two populations from evolving towards the same phenotype (Price *et al.* 2014).

To model niche filling, we first considered the migration function given in Eq. (11) with *L* = 0, and we assumed that the diverging optimum functions ***θ***^(*n*)^(*t*) are regularly “filling” the interval [*−A, A*] for some constant *A* ≥ 0, over successive epochs of fixed length *T*; that is, for 0 ≤ *t* ≤ *T*,

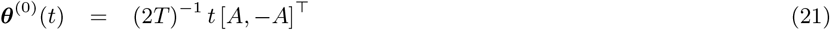

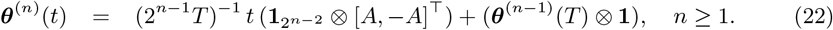

We refer to the left panel of Fig. 4 for a representation of the optimum functions over the first five epochs. In this particular example, if the migration function is the same for each species (Eq. (11)), the mean disparity 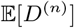 converges to a limiting value as *n* → ∞, given by

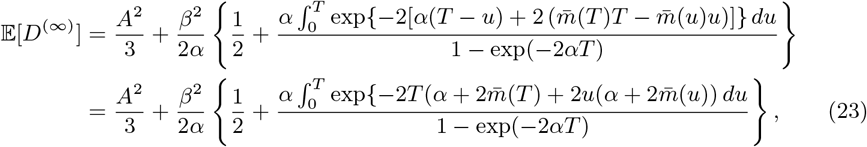

where 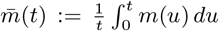; see Proposition SI-D.1. Note that we slightly abuse notation here, because we use 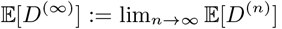. The term *A*^2^/3 in Eq. (23) corresponds to the variance of a uniform random variable in [−*A*, *A*], and the term *β*^2^/(2*α*) corresponds to the asymptotic variance of an OU process with no migration. The factor in the curly bracket accounts for migration (it reduces to 1 when there is no migration).

**Figure 2:**
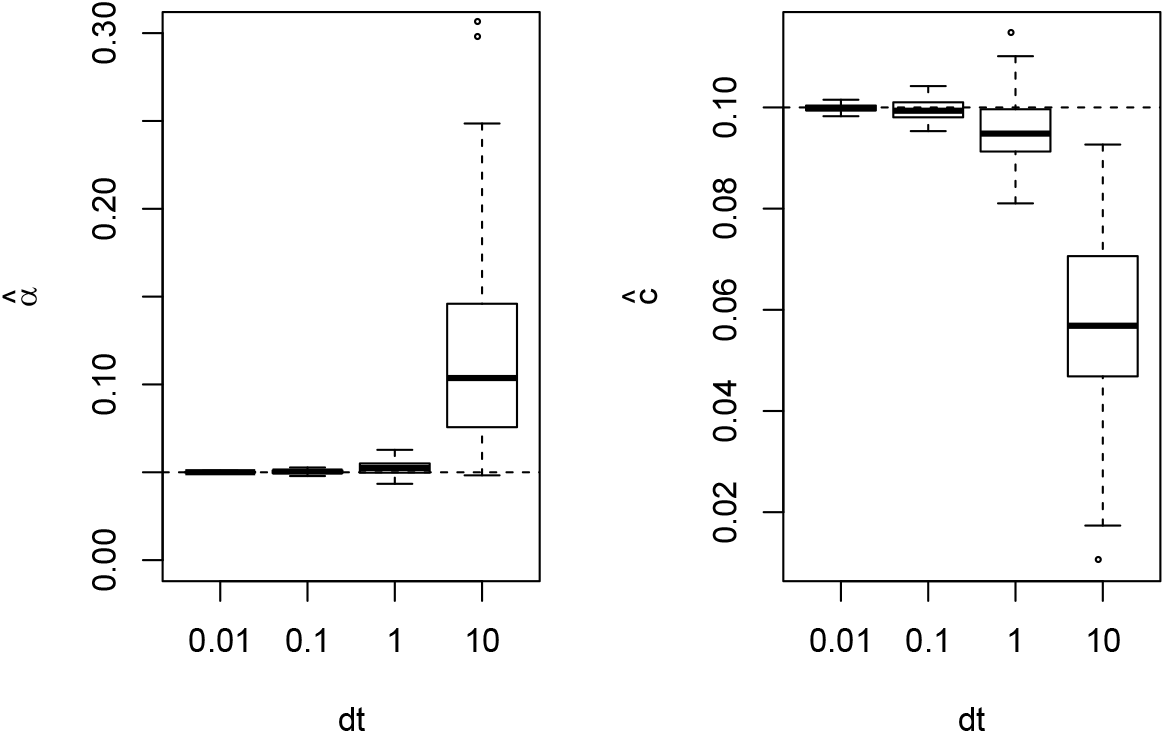
Empirical properties of the estimators for the selection coefficient *α* (left) and the migration parameter *c* (right). Each boxplot represents parameter estimations from 100 simulated phenotypic trajectories for two subpopulations under an OU process with *β* = 0.01 and four different step sizes *dt*.

**Figure 3:**
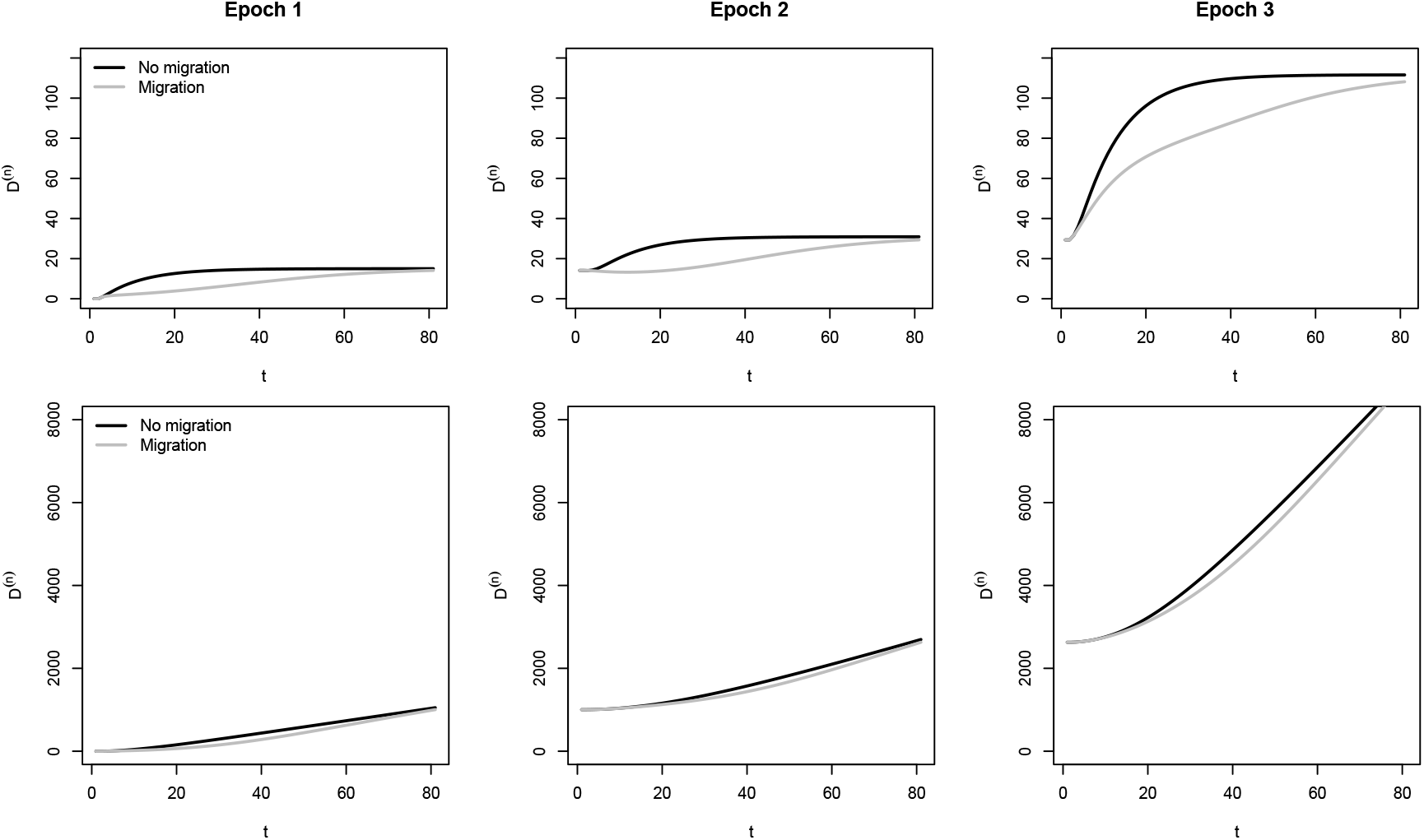
Phenotypic disparity *D*^(*n*)^ (Eq. (15)) for three consecutive epochs. Upper panels: disparity for stabilising optima 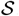. Lower panels: disparity for diverging optima 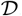. For scenarios with migration, we use *c*_1_ = 0.075 in Eq. (11) with *L* = 0. Optima per epoch were taken from Table SI-2. *β* was set to 0.

**Figure 4:**
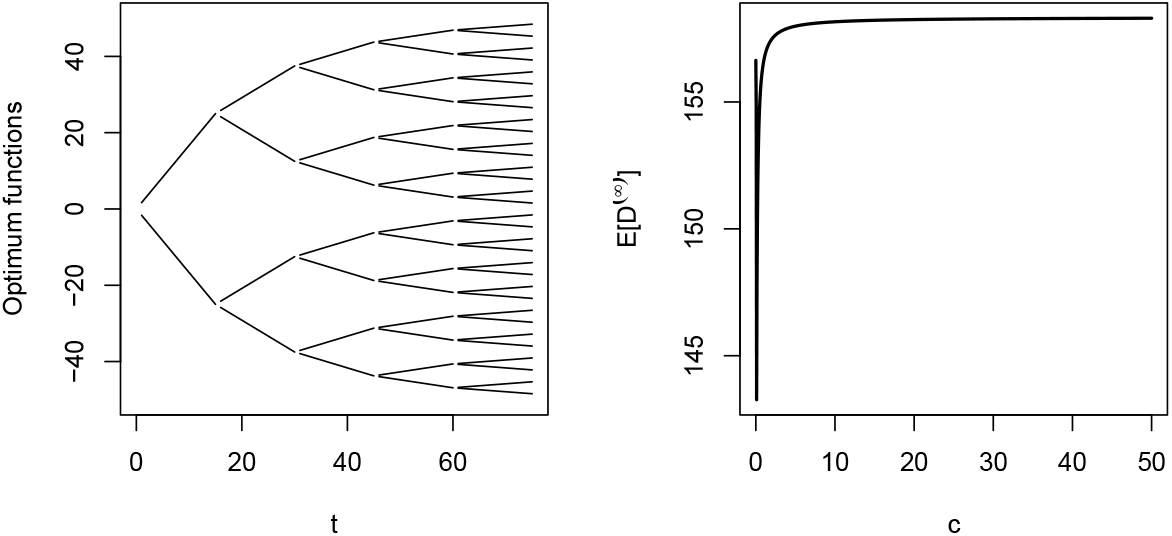
Left: Optimum functions as given by Equation (22) with *A* = 50 for *n* = 1, …, 5 and *m*(*t*) = 0.5 exp(*−ct*), where *c* is fixed such that *T* = 15. Right: asymptotic mean disparity as a function of the migration parameter *c*.

Next, we considered the case where the migration function depends on the differentiation function (Eq. (12)). If the slopes of the optimum functions are kept the same as in the previous case, during the *n*th epoch, the differentiation function then takes the form *d*^(*n*)^(*t*) = *At*/(2^*n*−1^*T*), and the length of the *n*th epoch is

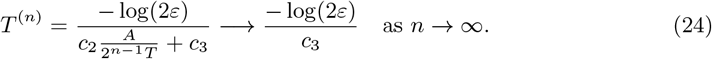

Hence, the effect of differentiation disappears asymptotically. In this case, the optimum functions are not confined within the interval [*−A, A*] (see top left of Fig. 5). The maximum absolute value of the optimum functions after *n* epochs is given by

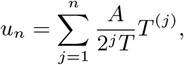

whose limit is finite and given by

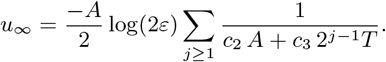

Note that this value can be larger than *A* (see top left of Fig. 5 where it is already above 120 after five epochs while *A* = 50). It is much harder to characterize the asymptotic behaviour of the mean disparity in this setting.

**Figure 5:**
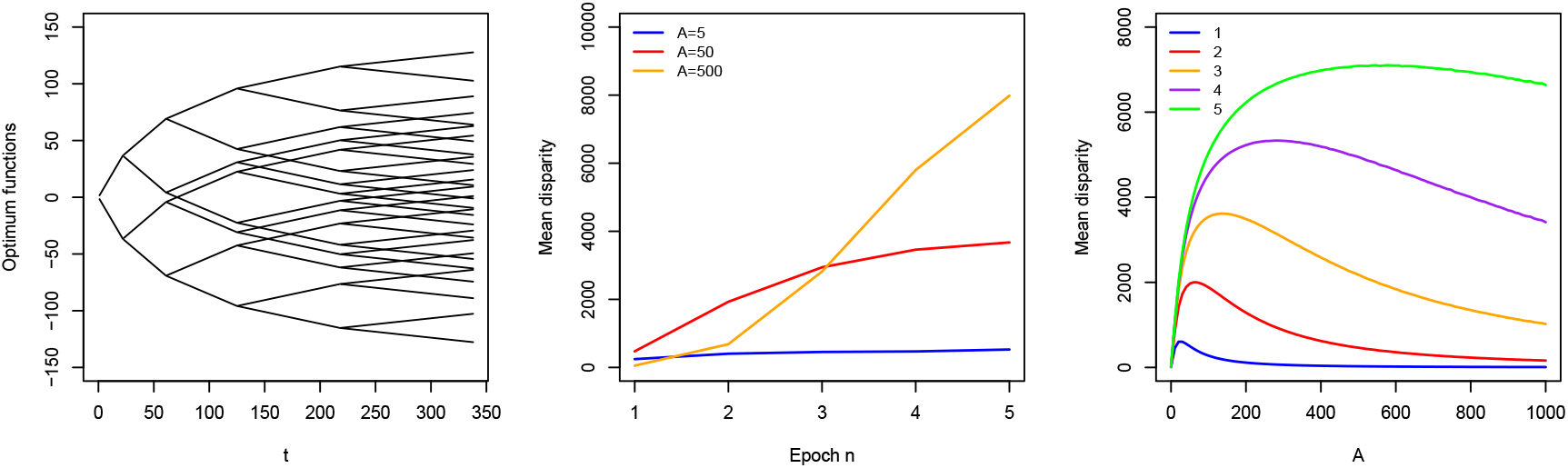
Left: Optimum functions as given by Eq. (22) with *A* = 50 for *n* = 1, …, 5. and *m*(*t*) = 0.5 exp(−0.1*d*(*t*)− 0.05*t*), where *d*(*t*) is the differentiation function. Center: mean disparity over five epochs for three values of *A*. Right: mean disparity at the end of five epochs (labeled 1 through 5), as a function of *A*.

## 3 Results

Following the same scheme as above we describe the results in three sections: results for the model, the estimators, and for the case study of niche filling. We give special attention to the results on a single epoch, since they can be directly translated to multiple ones.

### 3.1 Biological model

Without migration between the two subpopulations, the mean phenotype of each subpopulation reaches the optimum at a speed dictated by the selection coefficient *α* and a strong *α* will result in a fast convergence of 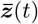 to ***θ***(*t*) (Fig. 1a). In the presence of migration, however, the speed at which the optimum is reached is slower (Fig. 1b) and different combinations of selection and migration will counteract each other to determine the speed at which the optima will be reached (Fig. 1c). When *β* > 0, stochastic fluctuations alter the path of 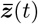, but the overall trend remains (Fig. 1d). For a period of time *L* of total mixing, the two subpopulations behave similarly and they remain together even when the two optima differ greatly (see Eq. (11)). However, as soon as *m*(*t*) starts decreasing (after time *L* = 500 in this example), the optima will be reached once again (Fig. 1e). Finally, if *m*(*t*) depends also on the distance *d*(*t*) between *θ*_1_ and *θ*_2_ (Eq. (12)), then the initial approach to the optimum can be faster than in the case where *m*(*t*) does not depend on *d*(*t*) (Fig. 1f versus 1b).

### 3.2 Estimators

#### 3.2.1 Joint estimation of *α* and *m*(*t*)

To validate our estimator 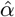 and the estimator of the migration parameter *ĉ* (*c* = *c*_1_ in Eq. (11) with *L* = 0) along one epoch, we simulated 100 population trajectories following an OU process with fixed *α* = 0.05, *c* = 0.10, and *β* = 0.01 for one epoch of fixed time *T* and various step sizes *dt*. We used Eq. (19) to compare the estimated *α* with the true value used in the simulation. Likewise, we compared the numerical solution for *c* with the true value used for simulations. The accuracy of the parameter estimates is directly related to the number of sampling points taken from the population trajectories, that is, inversely proportional to *dt* (Fig. 2). In other words, since *T* is fixed, a smaller *dt* results in more sampling points, thus increasing the accuracy of the estimation.

We also validated the estimators of *α* and *c* using the algorithm described in section SI-G, which generates the distribution of phenotypic points rather than the mean phenotype. With no data available from a second subpopulation, we can use Eq. (20) to estimate *α*. We, however, risk to underestimate *α* if there is ongoing migration from an unseen subpopulation. Consider the example in Figure SI-H.2a, where both trajectories were simulated using *α* = 0.05, but the “purple” trajectory experienced migration from an unseen subpopulation. When estimating *α* using Eq. (20), we observe that the “black” subpopulation has a correct *α* estimation, whereas we underestimate *α* for the “purple” subpopulation (Fig. SI-H.2b).

Therefore, to disentangle the effects of selection and migration when data from a single subpopulation is available, we need to look at the actual distribution of phenotypes within the population rather than at the simple mean. For instance, the distribution of phenotypes leading to the “purple” trajectory is, in fact, bimodal (Fig. SI-H.2c). On one hand, we have the bulk of the distribution that follows exactly the path of one subpopulation without migration, while we also see individuals that migrated from a second subpopulation petering out gradually as time passes by. The *α* parameter will be correctly estimated if the data from the bulk of the distribution is used, and this estimate will be well approximated even if only a couple of time points around the convergence value are considered (Fig. SI-H.2b, blue points). However, the robustness of 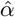 decreases as the two optima become closer, since it becomes more difficult to tell apart the two subpopulations (Fig. SI-H.3a).

Finally, by taking the difference between the number of individuals corresponding to both parts of the bimodal distribution shown in Figure SI-H.2c, we can approximate the migration function *m*(*t*) with high accuracy, as long as the optima of the two subpopulations are visibly different from one another (Fig. SI-H.2d, black points: actual *m*(*t*), red points: estimated *m*(*t*)). When the two optima are more similar, the accuracy of the estimation of *m*(*t*) decreases (Fig. SI-H.3b).

#### 3.2.2 Phenotypic disparity across epochs

We developed an estimator for the phenotypic disparity *D*^(*n*)^ while taking migration into account (Eq. (16)). Using the optima values shown in Table 2, we simulated the behaviour of *D*^(*n*)^ under cases with and without migration for three consecutive epochs. We found that disparity is reduced when migration is present and that this difference is bigger towards the beginning of an epoch (Fig. 3). The value of *D*^(*n*)^ will however become identical when migration vanishes. This behaviour is consistent for both types of optima: a stabilising optimum *S* (Fig. 3 upper panels) and a diverging optimum 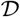 (Fig. 3 lower panels).

**Table 2:** **Experiment 1**: Parameters in the mixing vectors *m*^(*n*)^(*t*) and optimum vectors ***θ***^(*n*)^(*t*) corresponding to the first three epochs of a phylogenetic tree. In ***θ***^(*n*)^(*t*), 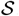 means *stabilising* (form (9)), and 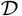 means *diverging* (form (10)).

| | $n = 0$ | $n = 1$ | $n = 2$ |
| --- | --- | --- | --- |
| $\mathbf{m}^{(n)}(t)$ | $c = 0.75$ | $c = 0.4$<br>$c = 0.55$ | $c = 0.2$<br>$c = 0.3$<br>$c = 0.4$<br>$c = 0.5$ |
| $\boldsymbol{\theta}^{(n)}(t)$ | $\mathcal{S}, a_1 = 10$<br>$\mathcal{D}, a_2 = -20$ | $\mathcal{S}, a_1 = 50$<br>$\mathcal{D}, a_2 = -10$<br>$\mathcal{D}, a_3 = 25$<br>$\mathcal{S}, a_4 = -30$ | $\mathcal{D}, a_1 = 5$<br>$\mathcal{S}, a_2 = 100$<br>$\mathcal{S}, a_3 = -300$<br>$\mathcal{D}, a_4 = -5$<br>$\mathcal{D}, a_5 = 10$<br>$\mathcal{D}, a_6 = 0$<br>$\mathcal{S}, a_7 = -100$<br>$\mathcal{S}, a_8 = 0$ |

### 3.3 Case study: niche filling

We used our model to study the filling of an ecological niche when phenotypic evolution includes migration. Niche filling occurs when there is ecological competition for limited amount of resources and phenotypes of two competing species cannot evolve towards the same optimum value.

We modelled niche filling in an interval of phenotypic values [*−A, A*] over successive epochs of fixed length *T* following Eq. (21) and (22). When *A* = 0, we are in the particular case where ***θ***^(*n*)^(*t*) = **0** for all *n* ≥ 0 and *t* ≥ 0. When *A* = 50, we can already see the “filling” effect of a niche with optima ranging between −50 and 50 (Fig. 4 left). Over the epochs, the mean disparity increases to a limit given by Eq. (23). We investigated the role of the migration rate in determining this limit, assuming that *m*(*t*) decreases exponentially at a rate *c* (Fig. 4 right). We see that there is a sharp drop in the asymptotic mean disparity as *c* increases from 0, then followed by a slow increase towards the constant value *A*^2^/3 + *β*^2^/(2*α*) (value of 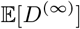 when *m*(*t*) ≡ 0 in Eq. (23)), indicating a larger mean asymptotic disparity when there is less mixing in the population. The interpretation of the drop is more difficult.

We also considered the case where the migration function depends on the differentiation function *d*(*t*) (Eq. (12)). The intervals of time between speciation events are thus not constant and depend on the epochs, but converge to a limit (Eq. (24)). The slopes of the optimum functions ***θ***^(*n*)^ were kept as in Eq. (22), and their initial values at the beginning of each new epoch are the continuation of their values at the end of the previous epoch. The graph of the functions ***θ***^(*n*)^ corresponding to *A* = 50 and *m*(*t*) = 0.5 exp(−0.1*d*(*t*) − 0.05*t*) is shown in the left panel of Figure 5. The central panel of that figure highlights for three values of *A* the logistic increase of the mean disparity as a function of the number of epochs *n*. We see again that there is always a plateau and the height of the plateau increases with *A*, but the limit is harder to characterize in this case. Finally, the mean disparity is not necessarily monotonically increasing with *A* when the migration function depends on the differentiation function *d*(*t*) (Fig. 5 right panel). Indeed, for each epoch *n* there is some threshold value of *A* such that the mean disparity increases for *A* less than this threshold due to the fact that the optima are more spread out, and it decreases for *A* larger than the threshold due to the fact that the differentiation becomes larger, and therefore the speciation times are smaller and there is less mixing before speciation.

## 4 Discussion

### Model

We introduced a model with migration between subpopulations (of a single species or a single branch on a phylogeny) that can be applied to macroevolution and can be represented as an OU process. This model combines features of the quantitative genetics model of Lande (1980a) (see also Lande (1976)) and Ronce & Kirkpatrick (2001), but allows for decreasing gene flow between subpopulations. A similar model but where gene flow stays constant has been proposed by Bulmer (1971), although he focused more on the fate of polymorphic alleles under those conditions. In our case, we introduced decreasing gene flow to capture speciation caused from subpopulations tending towards different phenotypic optima as a result of selection. We showed that, as expected, migration reduces the speed at which speciation takes place (e.g. Gavrilets 2004), and we highlighted the counteracting effect of migration on selection (see Fig. 1 for an illustration). Here, we showed that the interplay between migration and selection (e.g. Wright 1931; Ehrlich & Raven 1969; Stanley 1979; Charlesworth *et al.* 1982; Hartl *et al.* 1997; Ronce & Kirkpatrick 2001; Gavrilets 2004) is valid not only for one epoch but also for multiple ones and thus crucially extends to macroevolutionary levels. For instance, we show that the (macroevolutionary) selection coefficient *α* can be vastly underestimated if migration is overlooked, and migration slows down the speed at which phenotypic disparity among species is reached (Fig. 3). Migration has also recently been incorporated in the macroevolutionary model by Bartoszek *et al.* (2017), but there, constant migration was introduced between branches of a phylogeny (not within like here), and thus has no direct link to speciation.

### Micro and macroevolution

Bridging the micro and macroevolutionary scales has been a concern for evolutionary biologists since Darwin, and different ways of connecting these two levels have been proposed (Arnold *et al.* 2001; Reznick & Ricklefs 2009; Pennell & Harmon 2013). For instance, natural selection, which is a microevolutionary force, has itself an impact on macroevolutionary patterns. This can be seen in small isolated populations, which can experience fast phenotypic change at the microevolutionary scale if selection on the traits is strong (Lande 1980a), and this isolation can eventually turn into cladogenesis. Other microevolutionary forces, such as genetic drift and mutation, can also impact macroevolutionary patterns (Hansen & Martins 1996). For instance, under drift-mutation balance, the covariance between species phenotypes decreases with time and equates 2*G*_*m*_*t*_*z*_, with *G*_*m*_ being the mutation variance and *t*_*z*_ phylogenetic time (Hansen & Martins 1996), which is proportional to our Eq. (7). However, migration has not been considered yet. Gene flow can span over the two time scales and constitutes an important link between micro and macroevolution. Here, we propose that the two time scales are linked by the characteristic time *T*_*c*_, which defines the time over which the optima change in each subpopulation. A small *T*_*c*_ leads to changes occurring at a microevolutionary scale, while a large *T*_*c*_ indicates changes occurring at a macroevolutionary scale.

### One epoch

Our model assumes that a population (or species) consists of two subpopulations with initial random mixing. Each subpopulation then evolves towards distinct optima and the migration rate decreases until it becomes negligible. Speciation occurs when migration stops, leading to ecologically dependent reproductive isolation (EDRI, Hendry 2004). Selection against migrants contributes to EDRI, but also a number of other factors including: 1) reduced mating between individuals from both subpopulations (e.g. Higgie *et al.* 2000; Kirkpatrick 2001; Nosil *et al.* 2003); 2) habitat preferences (e.g. Rice 1984; Bush 1994; Via 1999); and 3) return to a specific breeding location (Hendry *et al.* 2004). Our results show that migration will slow down the speed at which the mean phenotype will reach an optimum value (Fig. 1b,c). When individuals migrate into a subpopulation, the mean phenotype of the latter is pushed towards the phenotype of the new migrants (Fig. 1e). The effect on the population optimum is initially strong, but will decrease with the reduced numbers of migrants over time. A similar conclusion was reached when the effect of migration rate on two connected populations of equal sizes was studied (Ronce & Kirkpatrick 2001).

### Multiple epochs

A similar pattern was observed across multiple epochs: the speed at which the different optima are reached depends on the antagonising effects of the selection coefficient *α* and the migration function *m*(*t*). Here, we showed that the underestimation of *α* also happens across multiple epochs if migration is not considered. We observed this while analyzing the mean disparity *D*^(*n*)^ along successive epochs while accounting for migration (Fig. 3). Recall that disparity is an important and widely-used statistic in ecology and phylogenetics (e.g. Harmon *et al.* 2003; O’Meara *et al.* 2006; Slater *et al.* 2010; Harmon *et al.* 2010; Lumbsch *et al.* 2010). The lag in *D*^(*n*)^ due to migration happens because gene flow reduces the phenotypic variance, since the addition of migrants in the subpopulations will reduce the differences between the subpopulation mean phenotypes.

### Estimation of *α* and migration

Inferring selection and migration has been a primary interest of evolutionary biologists (e.g. Lynch 1993; Kingsolver *et al.* 2001; Aitken *et al.* 2008). Our approach to inferring the selection coefficient *α* at the macroevolutionary scale takes the phenotype evolutionary trajectory into account, and the corresponding estimator 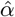 will be accurate around a time window of a couple tens of generations before and after the optimum value has been reached (and this approach can also be applied at the microevolutionary scale, to infer *α*_*m*_, Eq. (1)). After that, as we let the phenotypes evolve continuously around the same optimum for more generations 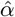 will slowly decrease (Fig. SI-H.2). This means that having phenotypic data sampled before or around the optimum is ideal when estimating selection. In any case, the estimator is robust even in cases when only a few data points have been sampled along the trajectory and as few as six time points leads to an accurate 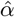 (Fig. SI-H.2b).

The migration function is more difficult to estimate unless one has time-series data. Such estimation is possible either if data from two subpopulations is available (Eq. (19)), or from one subpopulation only (Eq. (20)). Estimating migration with data from one subpopulation is possible because migrating individuals often display different phenotypic values if the two subpopulations have been already diverging. However, the difference in phenotypic optima between the two subpopulations has to be large enough to detect any effects. If the optima are close or similar to one another, an empirical estimation of the migration function becomes more difficult (Fig. SI-H.3). Current methods for estimating selection and migration jointly apply exclusively to genetic data (Hey & Nielsen 2004; Hey 2006; Mathieson & McVean 2013), whereas methods for estimating the selection coefficient *α* in phenotypic data do not take the effects of migration into account (Hansen 1997). Here, we show the importance of accounting for this evolutionary force since failing to do so results in a significant underestimation of the actual selective strength.

### Advantages of OU processes in phenotypic evolution

By taking into account selection, OU processes capture the evolution of quantitative traits and niches in a more realistic way than other models, such as BM or Early Burst (EB) models (Münkemüller *et al.* 2015; Boucher *et al.* 2014) (but see Pennell *et al.* 2015). For this reason, OU processes are widely used in comparative methods and quantitative genetics (e.g. Lande 1979; Martins *et al.* 2002; Rohlfs *et al.* 2013; Pan *et al.* 2014; Serrano-Serrano *et al.* 2015; Bartoszek *et al.* 2017). Pan *et al.* (2014), for instance, showed that changes in leaf-litter decomposability in several woody plants are better modelled by an OU process rather than BM or EB models. Serrano-Serrano *et al.* (2015) modelled the evolution of floral traits in neotropical Gesneriaceae and found a much better fit for an OU model for several traits. Rohlfs *et al.* (2013) also modelled gene expression using an OU process, but this time including within-species variation, which lead to the development of methods to detect changes in trait means and variances under OU (e.g. Khabbazian *et al.* 2016; Kostikova *et al.* 2016).

Finally, a direct (mathematical) advantage of our microevolutionary model with selection is that it is an OU process. Namely, it involves a linear phenotypic transformation, which makes the re-scaling of time straightforward, keeping the entire structure of the model maintained when we go from the micro to the macroevolutionary time scale (compare Eq. (1) with Eq. (5)). Therefore, the selection coefficient at the microevolutionary scale *α*_*m*_ becomes a cumulative selection coefficient *α* at the macroevolutionary scale, which amalgamates the effects of selection over multiple generations. This generalization should be further studied to investigate if the properties of the OU model at the microevolutionary scale can be extrapolated across speciation events and towards macroevolutionary time scales.

### Conclusions

We developed an extension of the OU model for phenotypic evolution that incorporates migration between populations within species while estimating the evolution of a quantitative trait. We show that, as expected at the microevolutionary scale, migration counteracts selection when populations diverge towards different optima for the quantitative trait, but our model allows to extend these results across multiple speciation events. The effect of migration is, therefore, important even for modelling trait evolution at the macroevolutionary scale and not accounting for this process can have important consequences for the estimation of key parameters such as selection levels typically considered in macroevolution.

## Author’s contributions

NS and LL conceived the ideas. PD and SH developed the project. PD and SH led the writing of the manuscript with substantial input from NS and LL. All authors contributed critically to the drafts and gave final approval for publication.

## Data accessibility

No new biological data has been generated in this project.

## Supplementary Information

### A Notation

In this paper we make use of the Kronecker product between matrices, which is defined as follows: if **A** is an *m × n* matrix and **B** is a *p × q* matrix, then the Kronecker product **A** ⊗ **B** is the *mp × nq* block matrix defined by

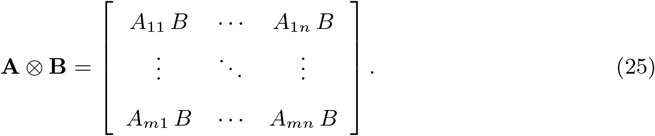

The symbol ^⊤^ denotes the matrix transposition. We let **1** := [1, 1]^⊤^, **J** := **1** *·* **1**^⊤^, *e* = [1, −1]^⊤^, and **E** = *e · e*^⊤^. We denote by **I** the identity matrix of size two, and we use the notation **1**_*x*_ for the column vector of 1’s of size *x*.

### B Trait evolution along one epoch

We first focus on the joint evolution of the phenotype of two subpopulations 1 and 2 forming one species between the birth of the species at time *t* = 0 until the next speciation event.

#### B.1 Dynamics along a single lineage

The system of stochastic differential equations characterizing the phenotypic evolution of the two subpopulations forming one species with common initial phenotype, 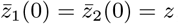, is given by

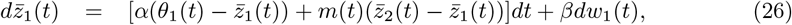

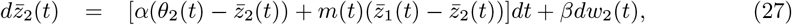

where *w*_1_(*t*), *w*_2_(*t*) are two independent Wiener processes, *α* denotes the strength of selection, and *β* describes the rate of stochastic evolution away from the optimum. Letting

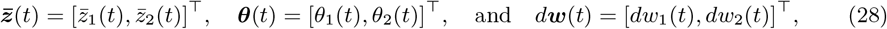

Eq. (26,27) may be rewritten in matrix form as

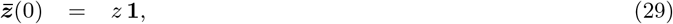

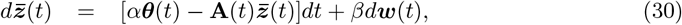

where

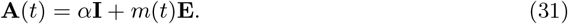

We let ***θ***(0) = *θ***1** for some constant parameter *θ*, because the optimum is initially the same for the two subpopulations forming a new species. Eq. (30) describes a multivariate inhomogeneous time-dependent Ornstein-Uhlenbeck (OU) process (see (Gardiner 2009, Section 4.5)).

We now solve (29,30) for *t* ∈ [0, *T*] following (Gardiner 2009, Sections 4.5.8 and 4.5.9). The homogeneous equation corresponding to (30) is the deterministic equation

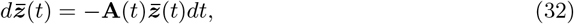

which is soluble since **A**(*t*)**A**(*u*) = **A**(*u*)**A**(*t*) for any *t, u* ≥ 0, and has the solution

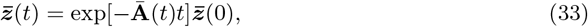

where **Ā**(*t*) is given by

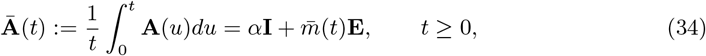

with

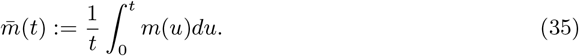

The general solution of (29,30) is given by

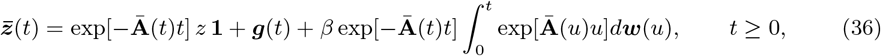

where ***g***(*t*) is the deterministic 2 × 1 vector

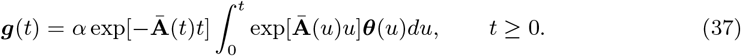

If *z* is deterministic or normally distributed, then 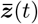 is normally distributed for any *t* ≥ 0.

Thanks to the particular form (34) of the matrix **Ā**(*t*), its exponential can be simplified, as we show in the next lemma.

##### Lemma B.1.

*For any t* ≥ 0,

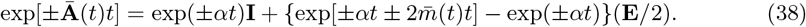

*In particular*, exp[−**Ā**(*t*)*t*]**1** = exp(*−αt*)**1**.

*Proof*. First observe that for *k* ≥ 1, **E**^*k*^ = 2^*k−*1^**E**. Then, using the binomial theorem for commuting matrices,

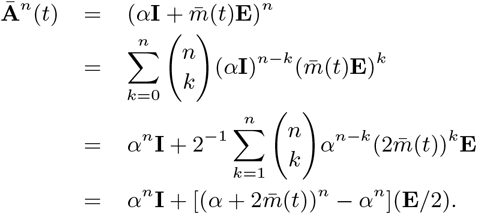

It follows that

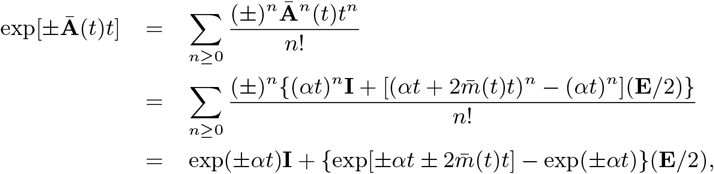

and since **E1** = **0**, we obtain the result.

As a consequence of Lemma B.1, the vector ***g***(*t*) can be rewritten as

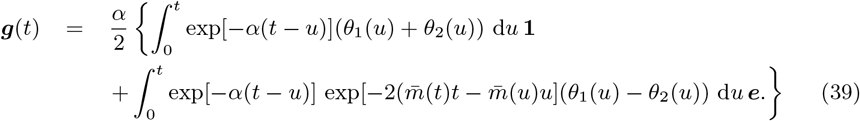

This expression can be further simplified if ***θ***(*u*) takes some special form. For instance,

- if ***θ***(*u*) = *θ*(*u*)**1**, that is, if the optimum functions are the same for the two subpopulations forming a species, then by (39) we obtain

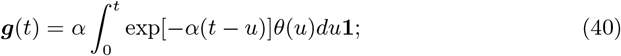
- if *θ*_1_(*u*) = *a* + *bu* and *θ*_2_(*u*) = *a − bu* (opposite linear functions with origin *a* and slope *b*), then *g*(*t*) simplifies to

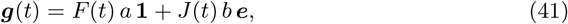

where

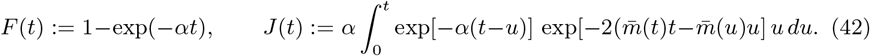

#### B.2 Dynamics of the mean and variance

We assume that the common phenotype *z* at time 0 is normally distributed with mean *μ* and variance *σ*^2^. We are now in a position to fully characterise the solution 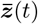 of (30).

##### Proposition B.2.

*For any time* 0 ≤ *t* ≤ *T, the random vector of mean phenotypes 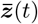 follows a multivariate normal distribution 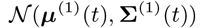*, *with* 2 × 1 *mean vector μ*^(1)^(*t*) *and* 2 × 2 *covariance matrix* Σ^(1)^(*t*) *given by*

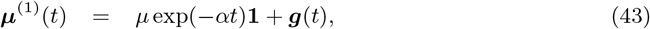

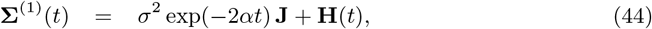

*where* ***g***(*t*) *is given by* (39), *and*

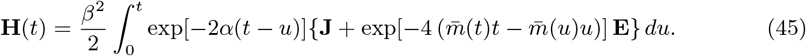

*Proof*. To obtain the expression for the mean, we take the expectation of the right-hand-side of (36), noting that 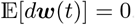. To obtain the covariance matrix, we take the expectation of 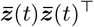 using (36) again, noting that **Ā**(*t*) = **Ā**^⊤^(*t*) and that 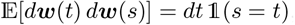, which leads to

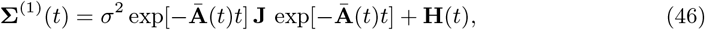

where

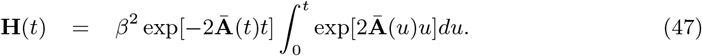

The final expressions (43) and (44) are then derived after some algebraic manipulations using Lemma B.1.

The first term in Σ^(1)^(*t*), *σ*^2^ exp(−2*αt*) **J**, takes into account the covariance induced by the common initial value *z* of 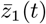 and 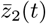, while the second term, **H**(*t*), takes into account the covariance induced by the Brownian noises acting on the two variables, and the mass exchange between the branches when *m*(*t*) *> ϵ*.

##### Remark B.3.

*In this setting we assumed that w*_1_(*t*) *and w*_2_(*t*) *are independent. The result can be generalized to the case where the two Wiener processes are not independent. In that case, we define ρ*(*t*) := Cov(*w*_1_(*t*), *w*_2_(*t*)), *and we can show that* Cov(*w*_1_(*t*), *w*_2_(*s*)) = *ρ*(min(*s, t*)) *for all s, t* ≥ 0. *The matrix* **H**(*t*) *then becomes*

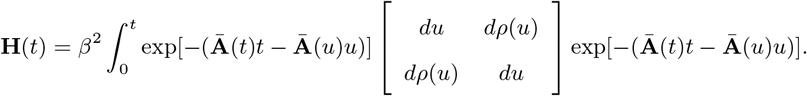

*In particular, if w*_1_(*t*) = *w*_2_(*t*), *then ρ*(*t*) = *t, and using Lemma B.1, we obtain*

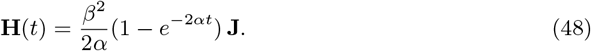

### C Trait evolution along the entire phylogenetic tree

We now consider the full process starting at time *t* = 0 with one species with mean phenotype *z ~ N*(*μ, σ*^2^) that splits into two subpopulations, and where migration occurs at rate *m*(*t*). In our model, each branch segment of the phylogenetic tree corresponds to two subpopulations evolving according to a two-dimensional OU process. There is a first speciation event at time *T* = inf{*t*: *m*(*t*) ≤ *ϵ*}, for a chosen value of *ϵ*. After that time, there is negligible mixing between the two subpopulations which give rise to two new species evolving independently of each other, *conditionally* on their initial mean phenotype 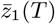 and 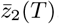. Again, each new species is made up of two subpopulations, and the process continues.

#### C.1 Constant mixing function *m*(*t*)

##### C.1.1 Dynamics of the mean and variance

If the migration function *m*(*t*) is deterministic and identical for all species, the next speciation events will happen at times 2*T*, 3*T*, etc. At time *nT* (corresponding to the end of the *n*th epoch), there will be exactly 2^*n*^ species in the process. The tree topology and branch lengths are therefore deterministic. However, the joint phenotypic distribution of the species at the end of each epoch is random and follows a multivariate normal distribution that we specify below.

For *n* ≥ 1, let 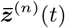, *t* ∈ [0, *T*], be the 2^*n*^−dimensional OU process describing the phenotypic evolution during the *n*th epoch, that is, between time (*n −* 1)*T* and time *nT*. We assume that we are given a sequence of (deterministic) functional vectors {***θ***^(*n*)^(*t*)}_*n*≥0_, of respective sizes 2^*n*^ × 1, defined for *t* ∈ [0, *T*], and containing the optimum functions corresponding to each epoch in the tree. That is, for *n* ≥ 1, ***θ***^(*n*)^(*t*) is the vector corresponding to 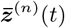. In order to ensure the continuity of the optimum function along each lineage, the vectors ***θ***^(*n*)^(*t*) must satisfy

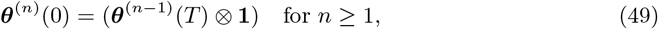

with ***θ***^(0)^ = *θ*.

In addition to the sequence of vectors {***θ***^(*n*)^(*t*)}_*n*≥1_, we define the related sequence of vectors {***g***^(*n*)^(*t*)}_*n*≥1_ of size 2^*n*^ × 1 as follows:

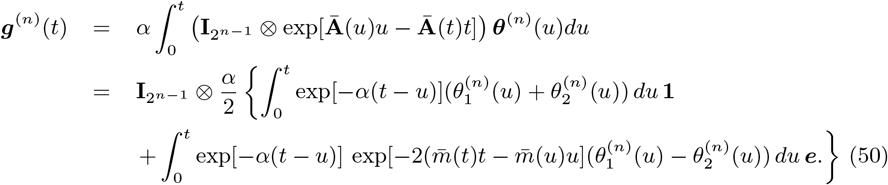

where **I**_2_*n−*1 denotes the identity matrix of size 2^*n−*1^.

Let 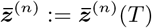 denote the random vector of phenotypes at the end of the *n*th epoch.

###### Proposition C.1.

*For n* ≥ 1, 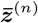 *follows a multivariate normal distribution* 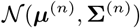 *of which the* 2^*n*^ × 1 *mean vector μ*^(*n*)^ *and the* 2^*n*^ × 2^*n*^ *covariance matrix* Σ^(*n*)^ *can be expressed recursively as*

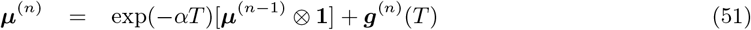

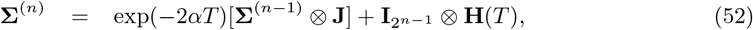

*with μ*^(0)^ = *μ and* Σ^(0)^ = *σ*^2^.

*Proof*. The recursion works by updating the initial (random) mean phenotype value of each species at the start of each epoch, which corresponds to the mean phenotype value of each subpopulation at the end of the previous epoch. The Kronecker products reflect the independent evolution of the mean phenotypes along each branch segment of the tree, conditional on their initial value.

###### Corollary C.2.

*For n* ≥ 1, *μ*^(*n*)^ *and* Σ^(*n*)^ *take the following explicit forms*

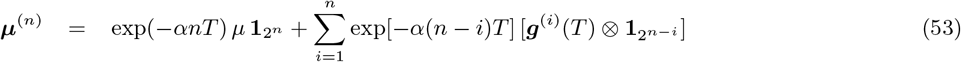

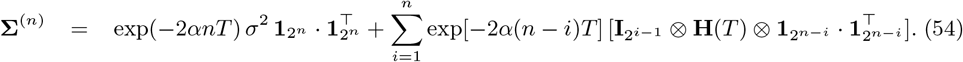

##### C.1.2 Evolution along a random lineage

Recall that each branch segment of the phylogenetic tree corresponds to two subpopulations evolving according to a two-dimensional OU process. One lineage of length *n* in the tree is thus one particular sequence of *n* branch segments controlled by a bivariate OU process, where at each branching point an optimum function *θ*_*i*_(*·*) (i.e. a direction) is chosen. Picking one lineage at random in a tree with *n* epochs is equivalent to selecting one of the 2^*n*^ leaves uniformly at random. The phenotype 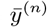 of this selected individual at time *nT* is given by

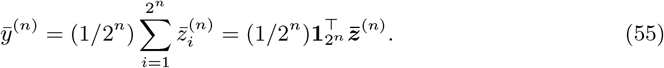

The random variable 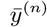 is thus simply the average mean phenotype at time *nT*, and is normally distributed with mean and variance

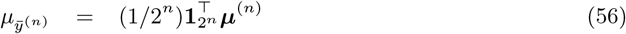

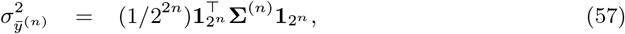

which satisfy a simple recursion: 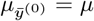 and 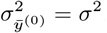, and for *n* ≥ 1,

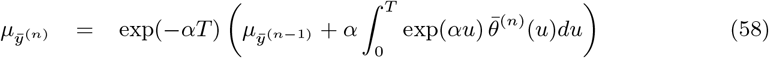

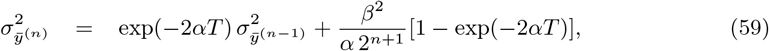

where 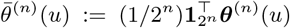 denotes the average optimum function during the *n*th epoch.

Asymptotically, as *n* → ∞, the variance vanishes, 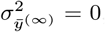, and 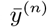 converges towards a constant

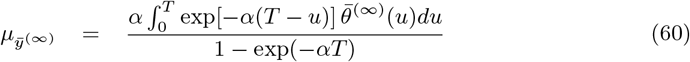

where 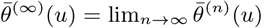.

##### C.1.3 Disparity of the phenotypic distribution

The disparity of of the multivariate vector 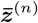, denoted by *D*^(*n*)^, is a scalar random variable which measures how much the mean phenotypes of the 2^*n*^ species present at the end of the *n*th epoch differ from each other. We define it as

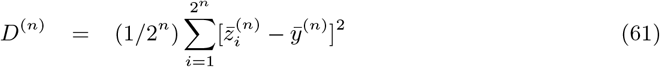

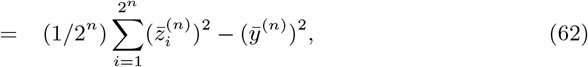

where 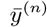 is given by (55). The disparity *D*^(*n*)^ is not to be confused with the variance of 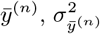, which measures the variability of the (random) average phenotype. The disparity corresponds to the sample variance of the mean phenotypes.

The first moment of *D*^(*n*)^ is given by

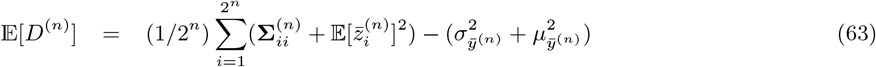

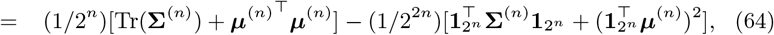

where Tr(Σ^(*n*)^) denotes the trace of the covariance matrix Σ^(*n*)^. In specific cases, such as the niche filling example, it is possible to characterise the asymptotic mean disparity 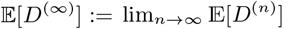, as we show in section D.

#### C.2 Variable mixing function *m*(*t*)

Here we assume that *m*(*t*) is deterministic but potentially different along each branch segment of the tree. That is, we define a sequence of functions {***m***^(*n*)^(*t*)}_*n*≥1_, where ***m***^(*n*)^(*t*) is a vector of size 2^*n−*1^ × 1 that contains the mixing functions corresponding to the 2^*n−*1^ systems of OU equations describing the phenotypic evolution during the *n*th epoch in the tree.

The definition of the sequence of mixing functions {***m***^(*n*)^(*t*)}_*n*≥1_ induces a sequence of corresponding speciation times {***T*** ^(*n*)^}_*n*≥1_ which are such that 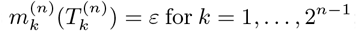; the branch segments of the tree can then have different lengths.

To each function 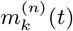 now corresponds a matrix

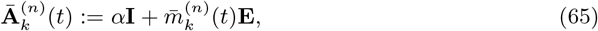

with

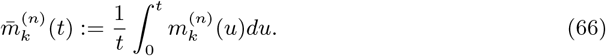

We define the related sequences of vectors {***g***^(*n*)^}_*n*≥1_ of size 2^*n*^ × 1 and matrices {**H**^(*n*)^}_*n*≥1_ of size 2^*n*^ × 2^*n*^ as follows:

- *g*^(*n*)^ contains 2^*n−*1^ block vectors of size 2 × 1, where the *k*th block vector, 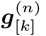, is defined as

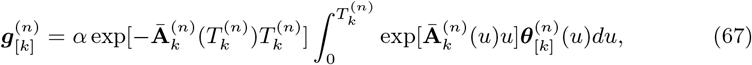

where 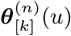 is the *k*th block-vector of size 2 × 1 in the 2^*n*^ × 1 vector ***θ***^(*n*)^(*u*), 1 ≤ *k* ≤ 2^*n−*1^;
- **H**^(*n*)^ is 2^(*n−*1)^ × 2^(*n−*1)^ block-diagonal, where the *k*th block matrix of size 2 × 2 on the diagonal, 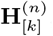, is defined as

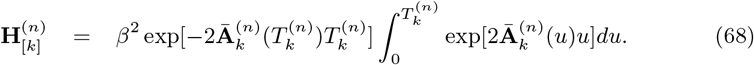

Like in (39) and (45), these expressions can be simplified using Lemma B.1. Let Diag[exp(*−αT* ^(*n*)^)] be the 2^*n−*1^ × 2^*n−*1^ diagonal matrix whose (*i, i*)th entry is 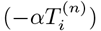. As before, 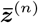 denotes the vector of phenotypes at the end of the *n*th epoch. Note however that species in the *n*th epoch may now be born at different time epochs. The random vector 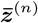 follows a multivariate normal distribution 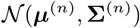 of which the mean vector and covariance matrix can be expressed recursively:

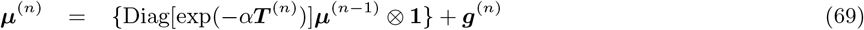

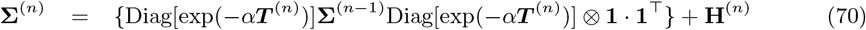

for *n* ≥ 1, with *μ*^(0)^ = *μ* and Σ^(0)^ = *σ*^2^.

Note that in the setting of variable mixing functions, it is not possible to obtain a recursive expression for the mean and variance of the average mean phenotype 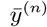.

### D Niche filling

In the niche filling example in the interval [*−A, A*] with fixed migration function *m*(*t*), the speciation time *T* is fixed. Let ***a***^(*n*)^ and ***b***^(*n*)^ denote the 2^*n−*1^ × 1 vectors containing the origins and slopes of the optimum functions ***θ***^(*n*)^, *n* ≥ 1. These vectors satisfy

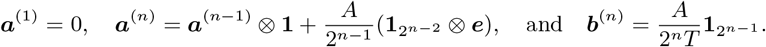

By (41) and (50), we then have ***g***^(*n*)^(*T*) = *F* (*T*)(***a***^(*n*)^ ⊗ **1**) + *J*(*T*)(***b***^(*n*)^ ⊗ ***e***), where *F* (*t*) and *J*(*t*) are defined in (42). Therefore, the mean vector and covariance matrix (53) and (54) of 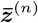 become

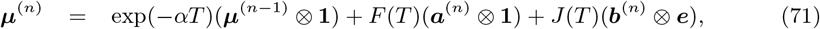

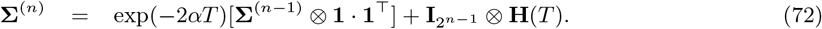

The asymptotic mean disparity takes a simple form, as we now show.

#### Proposition D.1.

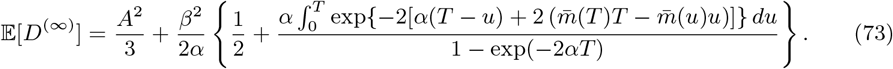

*Proof*. As *n* → ∞, due to symmetry, we have 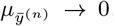, and we also have 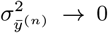. Therefore,

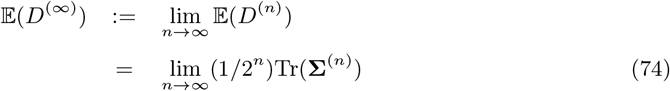

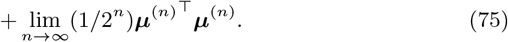

We first treat (74) and let *X* := lim_*n*→∞_(1/2^*n*^)Tr(Σ^(*n*)^). By (72), we have

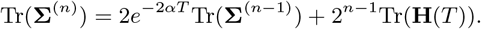

Dividing both sides by 2^*n*^, and taking *n* → ∞, we get *X* = *e*^−2*αT*^ *X* + Tr(**H**(*T*))/2, leading to

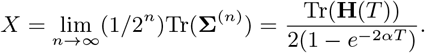

Next, we evaluate (75) and let *W* := lim_*n*→∞_(1/2^*n*^)*μ*^(*n*)^⊤^^*μ*^(*n*)^. Using (71), and the fact that **1**^⊤^**1** = ***e***^⊤^***e*** = 2, ***e***^⊤^**1** = **1**^⊤^***e*** = 0, we get

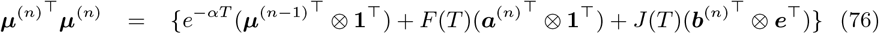

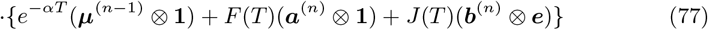

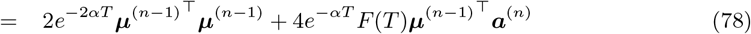

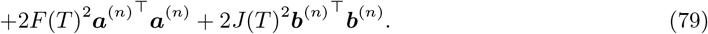

Dividing both sides by 2^*n*^, and taking *n* → ∞, we get

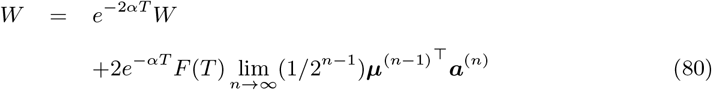

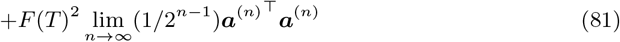

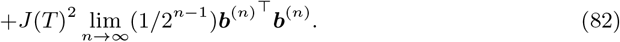

It remains to treat (80)–(82), which we do separately.

- Eq. (82): 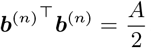, therefore (82) = 0.
- Eq. (81): Using the recursion for ***a***^(*n*)^,

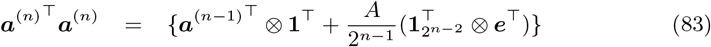

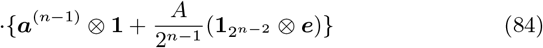

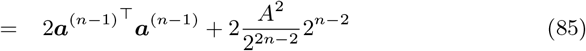

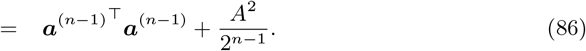 This is a first order recurrence equation with ***a***^(1)^⊤^^***a***^(1)^ = 0 whose solution is

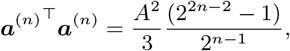

so we have

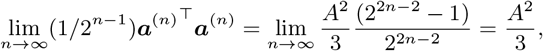

therefore (81) = *F* (*T*)^2^*A*^2^/3.
- Eq. (80):

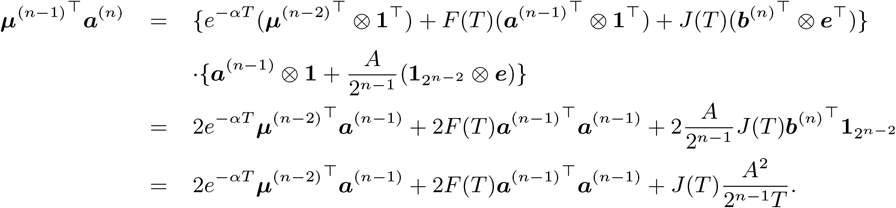 Let *Z* := lim_*n*→∞_(1/2^*n−*1^)*μ*^(*n−*1)^⊤^^***a***^(*n*)^. Dividing both sides by 2^*n−*1^, and taking *n* → ∞, we get *Z* = *e*^*−αT*^ *Z* + *F* (*T*)*A*^2^/3, therefore

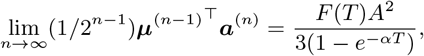

and

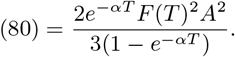

Coming back to the equation for *W*, we therefore have

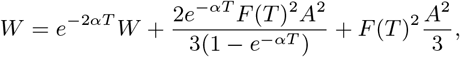

which, using the fact that *F* (*T*) = 1 *− e*^*−αT*^, simplifies to *W* = *e*^−2*αT*^ *W* + (*A*^2^/3)(1 *− e*^−2*αT*^), giving

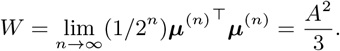

Summarizing, we have

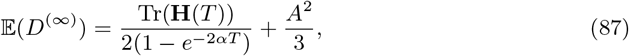

where,

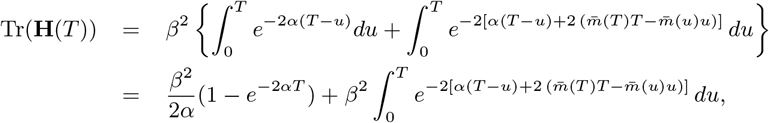

leading to (73).

### E Additional experiment

In this experiment, we fix *m*(*t*) = 0.5 exp(*−c*_1_*t*) with *c*_1_ = 0.5678 so that *T* = 15 (with *ϵ* = 10^−4^), and we consider the optimum function ***θ***^(*n*)^(*t*) depending on one parameter 0 ≤ *p* ≤ 5 as given in Table 3. These functions are illustrated in Figure H.4 for the extreme values *p* = 0 (Fig. H.4a) and *p* = 5 (Fig. H.4b). When *p* = 0 we are in the particular case where 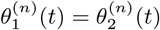 for all *n* ≥ 0 and *t* ≥ 0. The parameter *p* controls the percentage of increase in the parameter *a*_2_ of 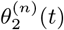 with respect to *a*_1_ in 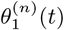. The mean disparity 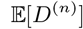 is plotted in Figure H.4c as a function of *p*. As expected, the mean disparity increases with *p*, but we also see that there is a threshold value *p*\* such that 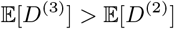 for *p* < *p*\*, and 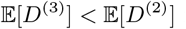 for *p* > *p*\*.

**Table 3:** Parameters in the optimum vectors ***θ***^(*n*)^(*t*) corresponding to the first three epochs of a phylogenetic tree. In ***θ***^(*n*)^(*t*), 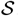 means *stabilising* (form (9)), and 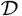 means *diverging* (form (10)).

| | $n = 0$ | $n = 1$ | $n = 2$ |
| --- | --- | --- | --- |
| $\boldsymbol{\theta}^{(n)}(t)$ | $\mathcal{D}, a_1 = 10$<br>$\mathcal{D}, a_2 = 10(1 + p)$ | $\mathcal{S}, a_1 = -10$<br>$\mathcal{S}, a_2 = -10(1 + p)$<br>$\mathcal{D}, a_3 = 30$<br>$\mathcal{D}, a_4 = 30(1 + p)$ | $\mathcal{D}, a_1 = 5$<br>$\mathcal{D}, a_2 = 5(1 + p)$<br>$\mathcal{S}, a_3 = -300$<br>$\mathcal{S}, a_4 = -300(1 + p)$<br>$\mathcal{D}, a_5 = -5$<br>$\mathcal{D}, a_6 = -5(1 + p)$<br>$\mathcal{S}, a_7 = 100$<br>$\mathcal{S}, a_8 = 100(1 + p)$ |

### F Estimation of the selection coefficient and the migration rate

As stated before, one of the main goals of this study is to analyze the bias on the estimation of *α* when failing to account for intraspecific migration. The motivation behind is that standard OU applications in macroevolution often aim at quantifying the amount of selection experienced by different species but they do not consider the effect that intraspecific gene flow has in these estimations. Therefore, given that in real phenotypic samples the selection coefficient *α* is unknown, we aim here at formulating two estimators: an estimator of *α* and an estimator of the migration rate parameter *c* = *c*_1_ in Eq. (11) when *L* = 0. Our model readily lends itself to derive such estimators by setting *β* = 0 in Eq. (5) and approximating these expressions as difference equations as follows:

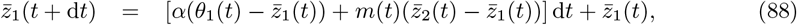

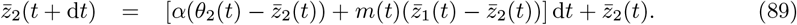

Following the same logic, in a second time step we have

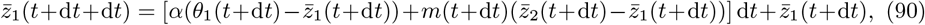

where the last term 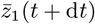 is given in in Eq. (88). So Eq. (90) becomes

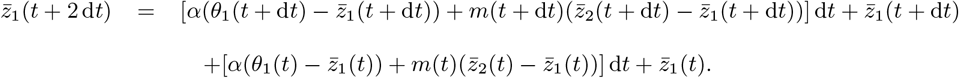

After *n* steps, we obtain

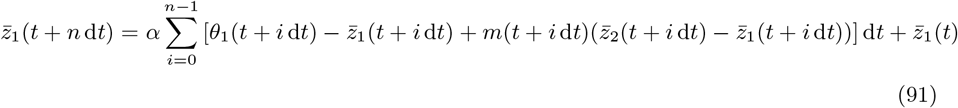

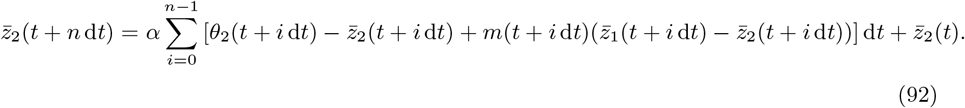

Adding Eq. (17) and (18) leads to

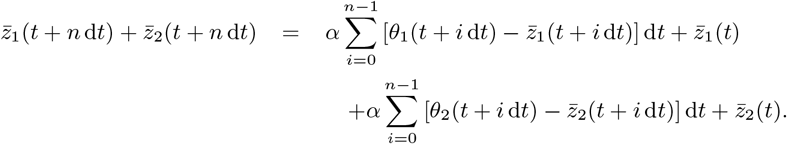

By rearranging terms, an estimator of *α* can be written in terms of the mean phenotype in the two subpopulations at times *t, t* + d*t, …, t* + *n* d*t* as

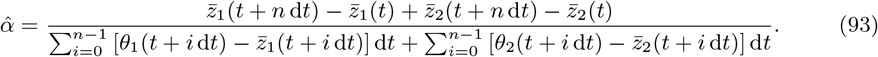

To obtain an estimator of the migration rate *c* we simply replace *α* in Eq. (17) or Eq. (18) with the value of Eq. (19) and solve numerically for *c*.

If there is no data on a possible second subpopulation from which migration could be taking place then *α* can be estimated from Eq. (19) using only the terms corresponding to subpopulation 1:

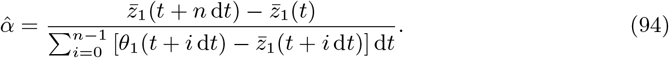

### G Simulations

This subsection describes a simulator that uses a standard OU process to generate phenotype trajectories with known *α* values. This simulator is used to 1) check the accuracy of 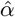, and 2) check the effect of gene flow in the estimation of *α* (accomplished by adding migrants from a second population).

#### Direct OU simulations

To check the accuracy of our estimator 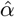 we simulated full population phenotype trajectories by using a standard OU process with the following steps:

1. With arbitrary values of 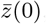, *α*, and *θ* (where *θ* is here a constant) we generated a trajectory of trait means 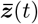 using equation 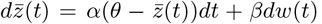. For the purpose of these simulations we set *β* = 0 and worked with the first term of the expression as a difference equation, as it is usually done when programming differential equations.
2. The vector of phenotype means 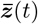 generated in step 1 for discrete values of *t* was then used to draw values from normal distributions with means 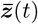 and some arbitrary variance to generate the corresponding population samples. This results in a population trajectory with a known *α*, a step that is necessary to validate the estimator described in Section F.

To check for the effect on 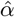 of migrants present from a different population we simulated a second population with the same parameters, varying only the optimum *θ*. The two populations then exchanged migrants at a rate given by Eq. (11) with *L* = 0. Notice here why it was necessary to simulate full population phenotypes and not only mean phenotypes, so that the two subpopulations can actually exchange migrants.

### H Supplementary figures

**Figure H.1:**
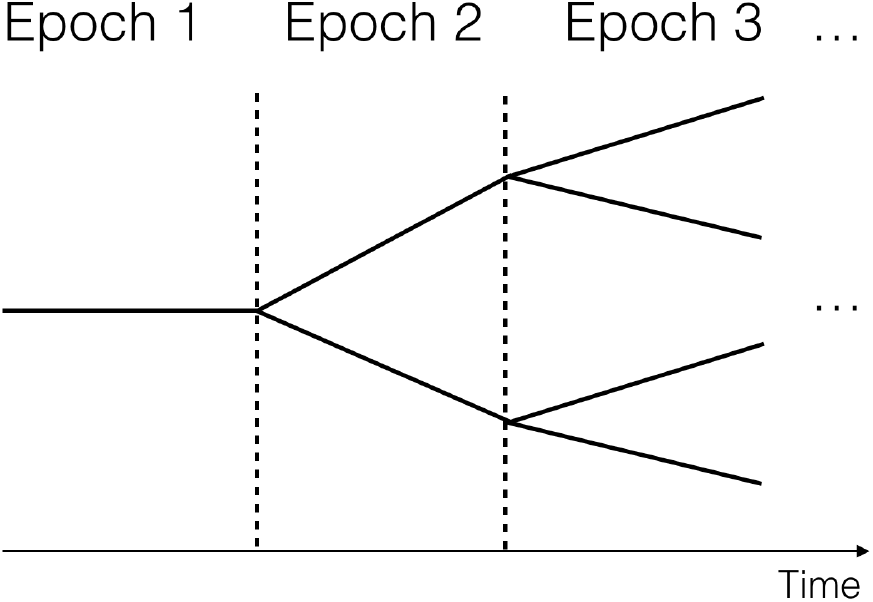
Schematic diagram of a phylogenetic (species) tree with three epochs. Here, there are 2^*n−*1^ coexisting species during epoch *n*, corresponding to 2^*n*^ subpopulations (*n* ≥ 0). Thus, in epoch *n* = 1 there are 2^0^ = 1 species, in epoch *n* = 2 there are 2^1^ = 2 species, and so on.

**Figure H.2:**
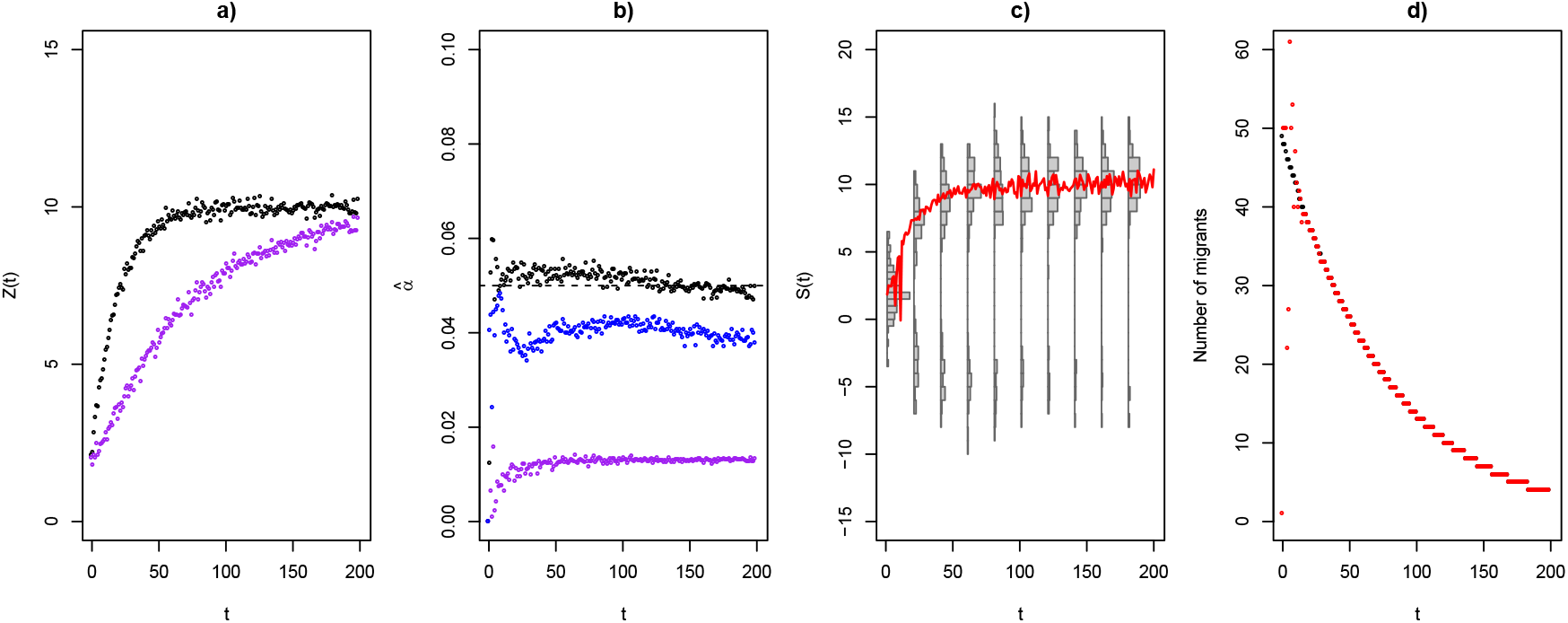
Estimation of *α* and empirical estimation of *m*(*t*). a) Trajectory of a mean phenotype *Z* following an OU process (simulated following the algorithm of section G) with *α* = 0.05 without incoming migration (black) and with migration from a second population (purple). b) *α* values estimated with Eq. (20) for the trajectory with migration (purple), without migration (black), and without migration but when data is sampled at only 6 time points around the convergence value (blue). c) Actual distributions of the “purple” phenotypes shown in panel a); the red line follows the mean phenotype along the distribution part with the highest density. d) True migration function of the trajectory shown with the purple circles in panel a) (black), versus the estimated migration function obtained from the distributions in panel c) (red).

**Figure H.3:**
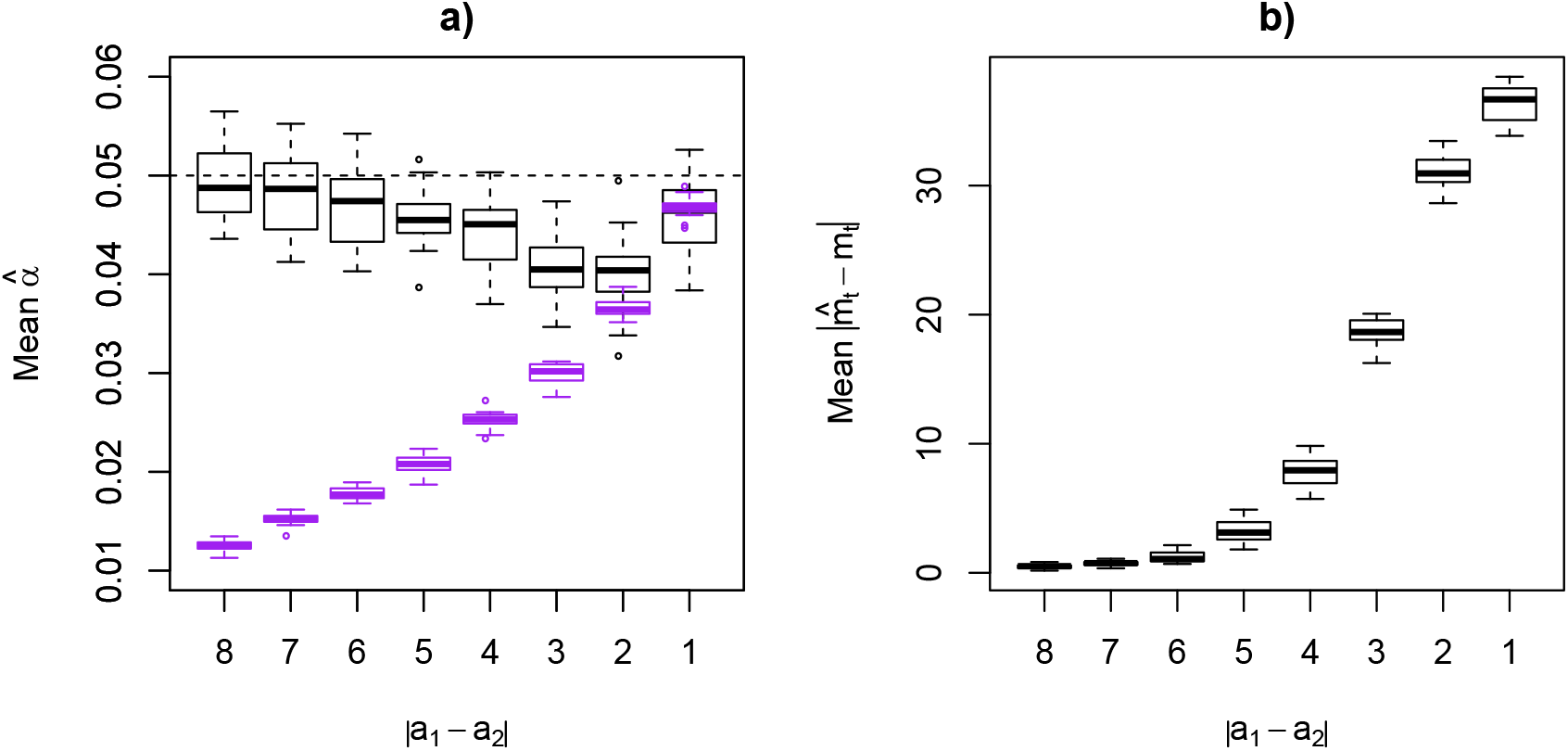
a) Robustness of 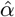 relative to the distance between the optima *a*_1_ and *a*_2_ of subpopulations 1 and 2, respectively. Black boxplots represent the mean of 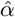 after 20 repetitions when accounting for migration. Purple boxplots represent the mean of 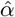 after 20 repetitions when not accounting for migration. The black horizontal dashed line shows the true *α* value. b) Robustness of 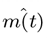 relative to the distance between the optima of subpopulations 1 and 2, measured as the mean absolute difference between the real an estimated migration function *m*(*t*) after 20 repetitions.

**Figure H.4:**
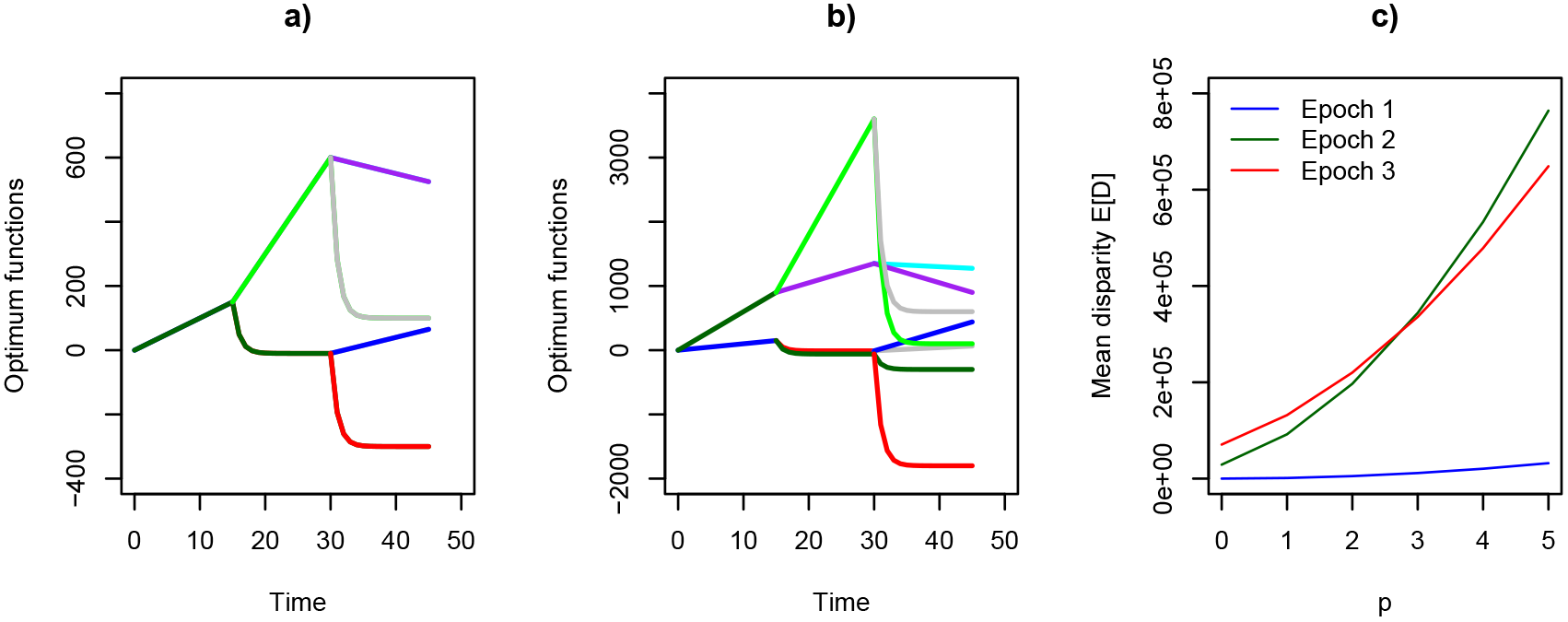
Optimum functions as given in table 3 for *p* = 0 (a), and *p* = 5 (b). c) Mean disparity *E*[*D*^(*n*)^] as a function of *p* for *n* = 3 epochs.

